# Live visualization of extracellular matrix dynamics during development and regeneration in zebrafish

**DOI:** 10.1101/2025.11.02.686082

**Authors:** Jingwen Shen, Ranjay Jayadev, Jianhong Ou, Ashley Rich, Kazunori Ando, Stefano Di Talia, David R. Sherwood, Kenneth D. Poss

## Abstract

Extracellular matrix (ECM) plays fundamental roles in animal development, regeneration, and disease. The difficulty of tagging endogenous matrix proteins in vertebrates has limited the understanding of ECM composition and dynamics in complex tissues. To visualize vertebrate ECM components, we tagged zebrafish Laminin, gamma 1 (Lamc1), Collagen, type I, alpha 2 (Col1a2), and Transforming growth factor, beta-induced (Tgfbi) using C-terminus in-fusion genome editing. Analysis of these knock-in lines revealed distinct expression of each protein in various tissues during development and regeneration. Fluorescent recover after photobleaching (FRAP) analysis further indicated that Lamc1 is stable in fin fold matrix but more dynamic in myoseptal matrix of developing zebrafish, while Col1a2 and Tgfbi are stable matrix components in myosepta. Strikingly, we found that Col1a2-mScarlet protein accumulates at the amputation plane during tailfin regeneration, where it remains concentrated for several days and distant from the regeneration blastema. This “foundation” region also displayed a distinct transcriptome suggesting active and dedicated events at the base of the regenerating appendage. Our resource enables live capture of ECM dynamics that can identify new events in developing and regenerating zebrafish.

**Summary statement:** Extracellular matrix resources for zebrafish development and tissue regeneration.

## INTRODUCTION

Extracellular matrix is composed of structural proteins, modification enzymes and signaling molecules, and is vital for development, organ formation and tissue regeneration. During early development, ECM components and ECM-bound growth factors provide cues for cell proliferation, differentiation, and assembly into complex structures. ECM also exerts key biophysical roles in different stages of tissue regeneration to enable healing and clearance of debris, tissue growth and remodeling (Chen et al., 2015; Wang et al., 2013).

Visualization of ECM components and their dynamic properties is crucial to understanding how ECM regulates diverse developmental events and regenerative processes. In vertebrates, direct ECM visualization in models like zebrafish has relied on ectopic overexpression of fluorescent protein-tagged ECM genes using a related regulatory sequence (Hino et al., 2024; Morris et al., 2018; Sánchez-Iranzo et al., 2018; Sharma et al., 2019). CRISPR-Cas9 based knock-in techniques facilitate tagging of the endogenous genomic locus, and studies in *C. elegans* have used in-frame fusion with fluorescent proteins to visualize 58 basement membrane (BM) proteins (Fan et al., 2020; Jayadev et al., 2019; Jayadev et al., 2022; Keeley et al., 2020; Naegeli et al., 2017; Srinivasan et al., 2025), a specialized form of matrix organized into thin and dense sheets of ECM surrounding most tissues (Feitosa et al., 2011). Studies using these endogenous knock-in strains have revealed surprising insights into differences molecules that stably associate with the basement membrane scaffolding versus dynamic BM components that rapidly mobilize on and off the basement membrane or diffuse within the scaffolding (Keeley et al., 2020).

The human core matrisome comprises ∼300 gene product contributors to ECM scaffolds (Naba, 2024). Collagens are the main ECM structural proteins, and 28 different collagen types exist in zebrafish. Non-collagenous structural ECM components include the crucial glycoproteins like laminin, which anchors basement membranes to cell surface receptors and exist in heterotrimers with different combination of α, β, γ subunits (Feitosa et al., 2011). Many matrisome-associated secreted factors (e.g. Tgfbi (Transforming growth factor, beta-induced)) reside in ECM and regulate diverse cellular events. Here, Here, we incorporated a flexible linker and tagged three ECM components in zebrafish by CRISPR-Cas9 mediated in-fusion knock-in -- laminin Lamc1, collagen Col1a2, and Tgfbi – and used live imaging to visualize localization, dynamics and turnover during development and tissue regeneration.

## RESULTS AND DISCUSSION

### A C-terminal knock-in strategy to establish an ECM toolkit

To tag endogenous ECM components in zebrafish, we used a strategy to introduce an eGFP or mScarlet fluorescent protein in frame at the C-terminus, as described (Levic et al., 2021). We designed sgRNAs to cleave a sequence in an intron, as well as a double-stranded DNA donor with part of the last intron and last exon followed by a flexible linker and sequences coding for a fluorescent protein (Fig. 1A, see Methods). Through coinjection of these components with NLS-Cas9, we tagged 3 different ECM components: the structural ECM protein Lamc1, the fibrous ECM component Col1a2, and the secreted signal protein Tgfbi which are essential for development and regeneration (Chen et al., 2015; Kim and Ingham, 2009; Metikala et al., 2021; Page et al., 2013; Parsons et al., 2002; Rajan et al., 2020; Sánchez-Iranzo et al., 2018; Sharma et al., 2019; Subramanian et al., 2018; Xia et al., 2022). These ECM proteins were chosen as representative ECM proteins based on distinct structures and functions and increased expression levels in different adult tissue regeneration contexts (Fig. S1A, B).

**Figure 1.**
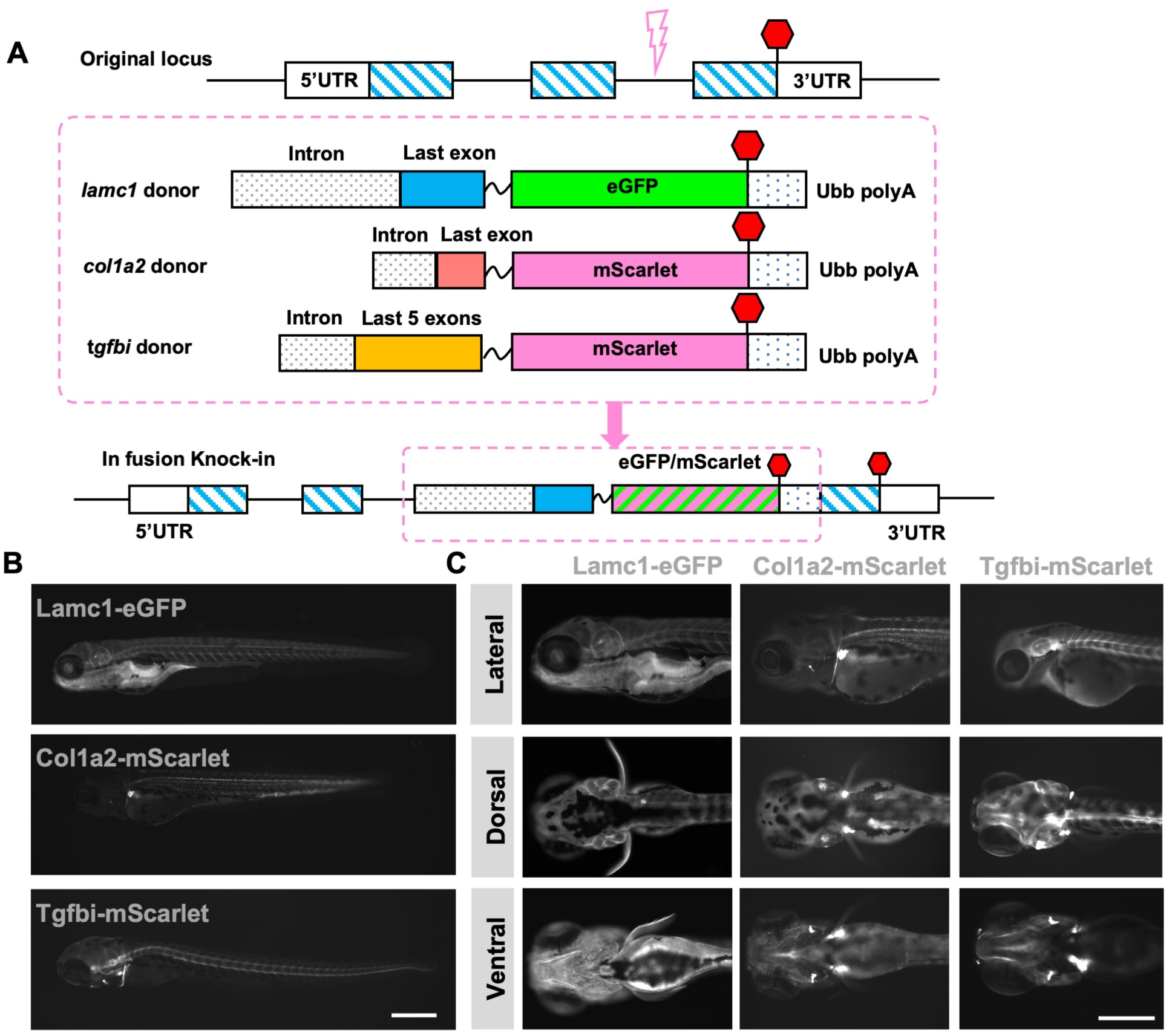
C terminal knock-in strategy to establish ECM knock-in resources. (A) C-terminal endogenous tagging strategy for 3 different ECM components. gRNA is designed to target sites in last introns for *lamc1* and *col1a2* genomic loci. For *tgfbi*, gRNA target site is located in the fifth intron. Partial intron and last exons were bridged by a flexible linker (GGSGGTGGSGGT) to fluorophore. Red hexagon, stop codon. Pink dashed box labels DNA donors for different target genes. (B) Images of *lamc1^eGFP^*, *col1a2^mScarlet^* and *tgfbi^mScarlet^* heterozygous larvae at 5 dpf. (C) Images of *lamc1^eGFP^*, *col1a2^mScarlet^* and *tgfbi^mScarlet^* heterozygous larvae from lateral, dorsal and ventral sides at 5 dpf. Scale bars are 500 µm (B, C).

Assessment of fluorescence signal from tagged proteins in embryos and larvae from stable knock-in lines indicated similar expression of each ECM fusion protein with its corresponding mRNA. The *lamc1* mRNA is present in transverse myoseptum, notochord, eye structures, fin fold, cardiac ventricle, and other structures (Parsons et al., 2002). We detected Lamc1-eGFP protein in *lamc1^egfp^* fish expressed throughout the whole body and enriched in the otic vesicle, pharyngeal arches, heart, pectoral fin bud, myoseptum, and notochord (Fig. 1B, C). *col1a2* transcripts are expressed in muscle, pectoral fin, branchial arches and cleithrum in zebrafish embryos and larvae, and transcriptional reporter lines have indicated similar expression domains (Rajan et al., 2020; Sharma et al., 2019) . In *col1a2^mScarlet^*animals, Col1a2-mScarlet was detected in cartilage of the lower jaw, cleithrum, and pectoral fin bud, and was especially enriched in joints and myosepta. Col1a2-mScarlet also formed punctate protein signals throughout the trunk (Fig. 1B, C). *tgfbi* mRNAs are known to be localized in splanchnocranium, vertebrae precursors, and pharyngeal arches in early stage zebrafish (Kim and Ingham, 2009). Tgfbi-mScarlet was prominent throughout *tgfbi^mScarlet^*somites and in the base of pectoral fin buds and myotome (Fig. 1B, C). We conclude that C-terminal knock-ins successfully labeled ECM components, and that the localization of tagged proteins is consistent with that of their mRNA transcripts.

### Larval protein dynamism differs between ECM components

ECM proteins are generally thought to be a stable fabric of tissue architecture, locked in place for long periods of time (Humphrey et al., 2014; Soans and Norden, 2021). Indeed, human matrix proteins are known to have long half-lives. In arteries, fibrillar collagen was estimated to perdure for 22 days, and elastin can last for many decades (Arribas et al., 2006). In adult mouse small intestine, pulse labeling of epithelial basement membrane components like laminin revealed turnover rates of weeks (Trier et al., 1990). In developing *Drosophila* embryos, turnover half-lives of perlecan and collagen IV from initiative induction stage at early embryogenesis were 7 and 10 hours, much faster than homeostatic status following development (Matsubayashi et al., 2020). In *Drosophila* tendon cells, tendon apical extracellular matrix (aECM) like proteinaceous matrix Dumpy (Dpy) undergoes filament-like transformation and showed fast recovery rates in minutes after photobleaching (Dong et al., 2014). Fluorescence recovery after photobleaching (FRAP) is a standard approach to understand the dynamics or kinetics (diffusion and on-off rates) of proteins like ECM at physiological protein concentrations. For instance, confocal FRAP measurements showed 2500 kDa aggrecan had a more rapid on-off association with the ECM compared to the 870 kDa hyaluronan in FITC-dextran solutes, because of aggrecan’s more compact and branched structure (Hardingham and Gribbon, 2000). Further, broadly endogenous mNeonGreen labeling of matrix proteins in *C. elegans* revealed static and dynamic ECM networks during morphogenesis and organ development (Keeley et al., 2020). From these studies, it is clear that ECM components can vary in their dynamics, and that mobilities of a given protein can be tissue- and stage dependent.

To assess ECM dynamics during embryogenesis, we performed confocal FRAP with all three knock-in lines generated. We first analyzed fluorescence recovery in the fin fold over a 10 min period, imaging every 10 s. Lamc1-eGFP is present in whole fin fold in a universally distributed pattern, whereas Col1a2-mScarlet is more punctate (Fig. 2A). Tgfbi-mScarlet was not detected in fin fold (Fig. 2A). In FRAP experiments, Lamc1-eGFP displayed little or recovery during the first 10 minutes after bleaching. Longer imaging times revealed less than 10% Lamc1-eGFP recovered after 4 h, with an average recovery half-life in fin fold of 11.3 ± 9.4 h (Fig. 2B-D). In contrast, the average recovery half-life of Col1a2-mScarlet in the fin fold was 4.1 ± 3.0 h (Fig. 2B). These results are consisting with previous finding that collagen and laminin are stable components in basement membrane (Keeley et al., 2020).

**Figure 2.**
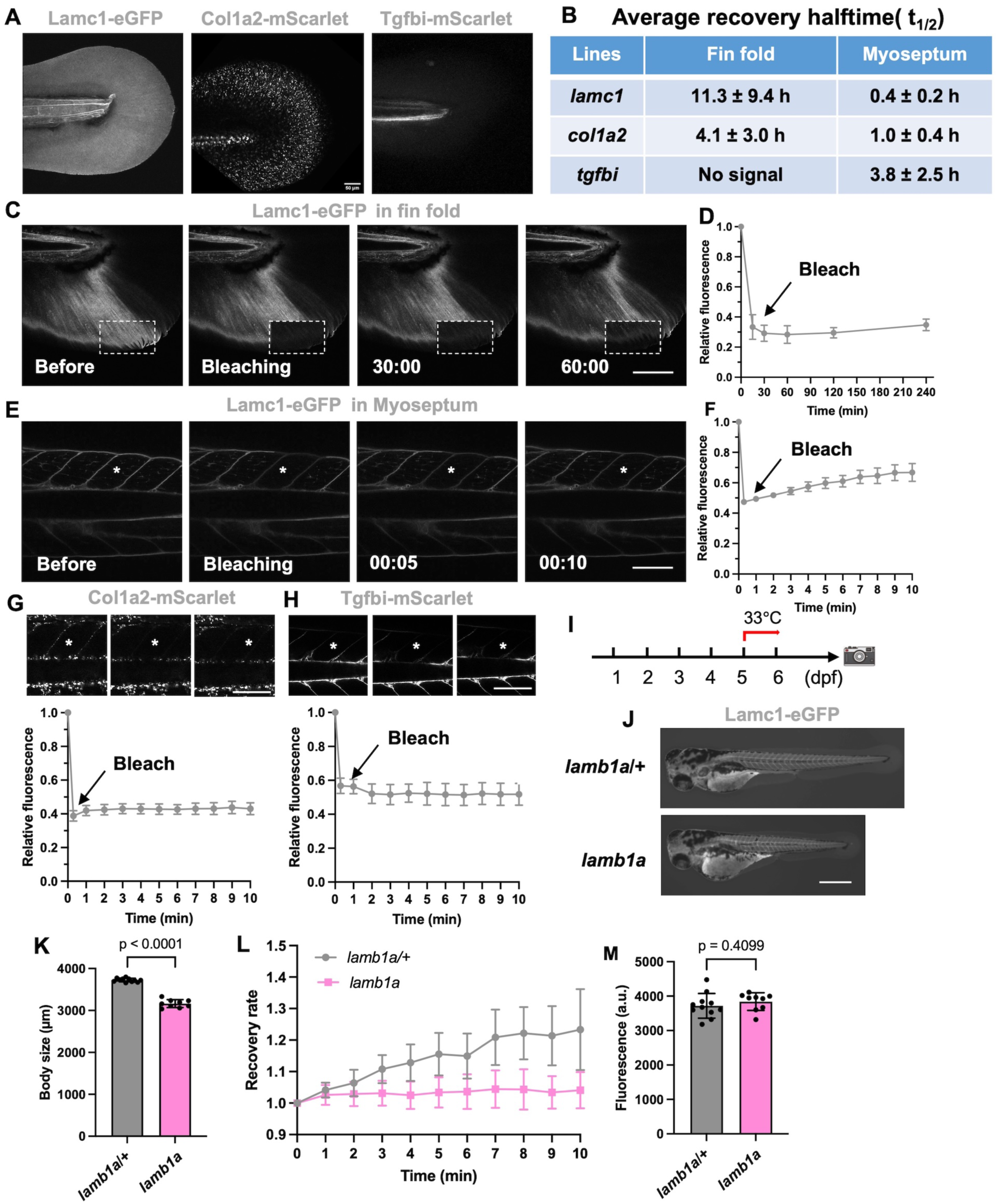
ECM component dynamism during embryogenesis. (A) Representative confocal images for Lamc1-eGFP, Col1a2-mScarlet, and Tgfbi-mScarlet before FRAP analysis. (B) Average recovery halftime for Lamc1-eGFP, Col1a2-mScarlet and Tgfbi-mScarlet in fin fold and myoseptum. Tgfbi-mScarlet was not expressed in fin fold at 5 dpf. (C and E) In vivo FRAP analysis of Lamc1-eGFP in fin fold (A) and myoseptum (C) at 5 dpf. Bleached area highlighted in the dashed box. Time: hour: minute. (n =6 and n=4 animals for each). (D and F) Line graph indicating quantified relative fluorescent recovery over time course in fin fold and myoseptum at 5 dpf. (G-H) FRAP analysis of Col1a2-mScarlet (n=8) and Tgfbi-mScarlet (n=7) in myoseptum at 5 dpf. Bleached area highlighted in the dashed box. Time: hour: minute. (I) Schematic for experiments with *lamb1a*; *lamc1^eGFP^* animals. Embryos with indicated genotypes were maintained at the permissive temperature (28.5 °C) until 5 dpf and shifted to the restrictive temperature (33°C) before analysis. (J) Representative images of *lamb1a*/+ and *lamb1a* larvae at 6 dpf in the *lamc1^eGFP^* background. (K) Quantification of body length for *lamb1a/+* and *lamb1a* larvae at 6 dpf in the *lamc1^eGFP^* background. (L) FRAP analysis of *lamb1a/+* and *lamb1a* larvae at 6 dpf in the *lamc1^eGFP^* background, assessing myoseptum. (n = 5 animals for each) (M) Relative fluorescence of myoseptum of *lamb1a/+* and *lamb1a* larvae at 6 dpf in the *lamc1^eGFP^* background. Scale bars are 50 µm (A), 100 µm (C,E,G,H), 500 µm (J)

Myosepta are specialized connective tissue sheet structures in somites that act as attachment points for muscle fibers, first visible at 1 dpf and continuing to mature throughout the lifespan (Keenan and Currie, 2019). They are like mammalian tendons and serve as transmitters of muscular contractility and are crucial for movement. Myosepta form from ECM components from 1-to-5 dpf and continue to maturation. We found that Lamc1, Col1a2 and Tgfbi fusion proteins are all enriched in myosepta, and thus suitable for FRAP experiments. FRAP experiments with *lamc1^eGFP^* revealed 20% recovery of signal just 10 min after photobleaching, and its recovery half-life of 0.4 ± 0.2 h was much faster than in fin fold (Fig. 2B, E, F). In contrast, Col1a2-mScarlet and Tgfbi-mScarlet showed no evidence of signal recovery at 10 min post-bleaching (Fig. 2G, H). These results indicate that Lamc1 is a dynamic matrix protein in myosepta but Col1a2 and Tgfbi are more stable.

To determine if the *lamc1^eGFP^* allele could help dissect dynamic properties of Lamc1 in development, we crossed temperature-sensitive mutations in *laminin beta 1a* (*lamb1a*) into this background (*lamb1a^pd110^*) (Chen et al., 2015). Laminins forms heterotrimers containing one alpha, one beta and one gamma chain (Timpl et al., 1979; Yurchenco and Cheng, 1993). To permit developmental progression and analyze Lamc1-eGFP dynamics in myosepta, we retained animals at permissive temperature for 5 days before shifting to the restrictive temperature of 33°C for 1 day, after which FRAP was performed (Fig. 2I). *lamb1a*; *lamc1^eGFP^* animals shifted to 33°C had shorter trunks than *lamb1a/+*; *lamc1^eGFP^* clutchmates, effects of *lamb1a* mutations (Fig. 2J, K). By performing photobleaching on *lamb1a*;*lamc1^eGFP^* embryos, we found Lamc1-eGFP in *lamb1a* mutant has a ∼20% lower recovery rate compared to *lamb1a/+* clutchmates (Fig. 2L), but its protein level was not changed in *lamb1a* mutant (Fig 2M). Together, these results suggest that the *lamb1a^pd110^* temperature-sensitive mutant does not affect levels or localization of Lamc1 to the basement membrane, but instead reduces its on and off association rate from the basement membrane.

### Visualization of ECM components across different adult organs

To investigate ECM-fluorescent protein fusion localization in adults, we examined the localization of several different tissues of *lamc1^eGFP^*, *col1a2^mScarlet^*, and *tgfbi^mScarlet^* zebrafish. In scales, bony discs of body armor that decorate the trunk surface, Lamc1-eGFP was present in the vasculature (Fig. 3E). These included vessels that run through the bony canals of the scale from base to periphery, as well as the less organized vessels penetrating the overlying skin (Fig. 3E). Col1a2-mScarlet was evident in the canals as well as punctate throughout the structure (Fig. 3E). In zebrafish kidney, which has filtering and hematopoietic functions, Col1a2- and Tgfbi-mScarlet marked proximal renal tubules (Fig. 3C). In cardiac ventricles, Lamc1-eGFP was evident at distinctly highly levels in the extracellular space surrounding muscle cells within the compact muscle layer, between the bilayer of epicardial cells, as compared to inner trabecular cardiomyocytes (Fig. 3A, Fig. S3A). Upon resection of the ventricular apex, a manipulation that stimulates vigorous regeneration (Poss et al., 2002), we noted increased expression in regenerating compact muscle. Col1a2-mScarlet expression was not detected above background autofluorescence in uninjured ventricles, but punctate signals were apparent at 14 dpa in the injury site adjacent to fibronectin deposits (Fig. 3B, Fig. S3B), resolving by 30 dpa. In uninjured spinal cord, all three knock-in products were not detectable by microscopy.

**Figure 3.**
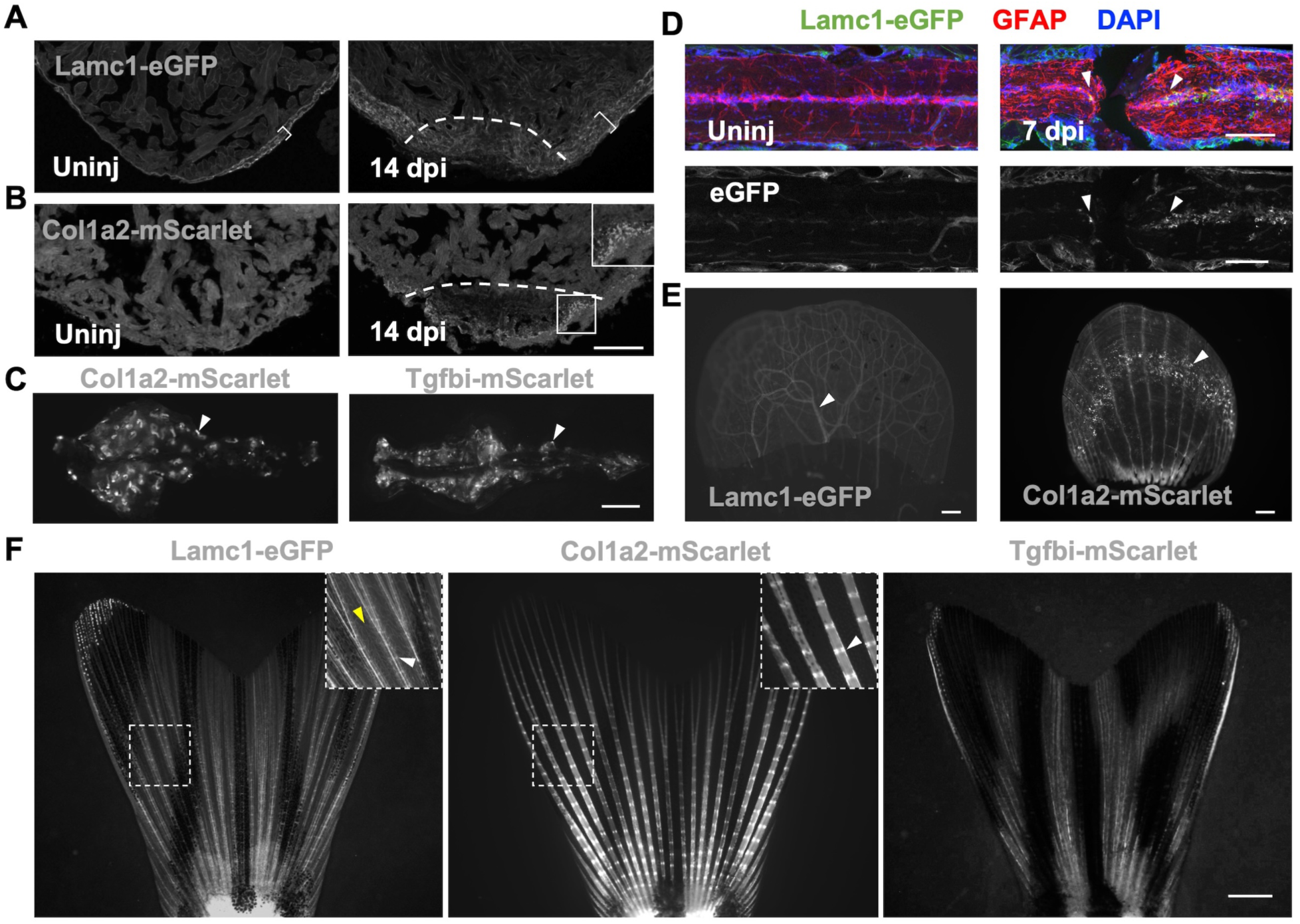
Localization and regenerative responses of different ECM components in adult zebrafish. (A) Confocal images of *lamc1^eGFP^* uninjured and 14 dpa ventricles. Dashed lines indicate approximate injury site; bracket indicates compact muscle layer. (B) Confocal images of *col1a2^mScarlet^*uninjured and 14 dpa ventricles. White box represents regions with Col1a2-mScarlet accumulation. (C) Col1a2-mScarlet and Tgfbi-mScarlet distributions in adult kidney. White arrowheads label renal tubules. (D) Lamc1-eGFP localization in uninjured and regenerating spinal cord (arrowheads; 7 dpi). (E) Lamc1-eGFP and Col1a2-mScarlet localization in a plucked scale. White arrowhead labels the Lamc1-GFP positive blood vessels (Left), and white arrowhead labels clustering signal formed by Col1a2-mScarlet (Right). (F) Representative images of Lamc1-eGFP, Col1a2-mScarlet and Tgfbi-mScarlet in adult caudal fins. White arrowheads, joint of fin rays. Yellow arrowhead, intraray vasculature. Scale bars are 100 µm (A, B, D), 1 mm (C, E, F).

However, Lamc1-eGFP was induced in cells of the central canal at both anterior and posterior stumps after a paralyzing complete transection injury at 7 dpi, which typically leads to regeneration within 6 weeks (Becker et al., 1998; Mokalled et al., 2016) (Fig. 3D). Thus, the resource we present here enables investigation of ECM components in a variety of adult organs, as uninjured tissues or in states of regeneration.

**Long-lasting accumulation of Col1a2 at the foundation of regenerating fins** Zebrafish fins are composed of several connected bony fin rays, each with repetitive segments connected by collagenous joints. Lamc1-eGFP in fins mainly localized to fin rays but was also found in blood vessels (Fig. 3F). Col1a2-mScarlet also localized to fin rays, especially at joints, while Tgfbi-mScarlet localization was similar to Col1a2-mScarlet, but present at lower levels (Fig. 3F).

To visualize the responses of 3 different ECM components in live regenerating fins, we performed caudal fin amputation and monitored regeneration at 4, 7, and 14 days post amputation (dpa). Lamc1-eGFP was present in the distal growing regions of regenerating rays at 4 dpa (Fig. S4A), remained distally throughout regeneration, though gradually decreasing in levels (Fig. 4A, C). In contrast, Col1a2-mScarlet was not detected before 4 dpa in the regenerating tissue (Fig. 4B, Fig. S4B). At 7 dpa, Col1a2-mScarlet was present at the amputation plane and continuing to the blastema, though especially enriched at the amputation site, where it was retained at 14 dpa and even as long as 60 dpa (Fig. 4B, C, Fig. S4C). Tgfbi-mScarlet was similar to Col1a2 and accumulated at amputation plane during fin regeneration (Fig. S4D, E).

**Figure 4.**
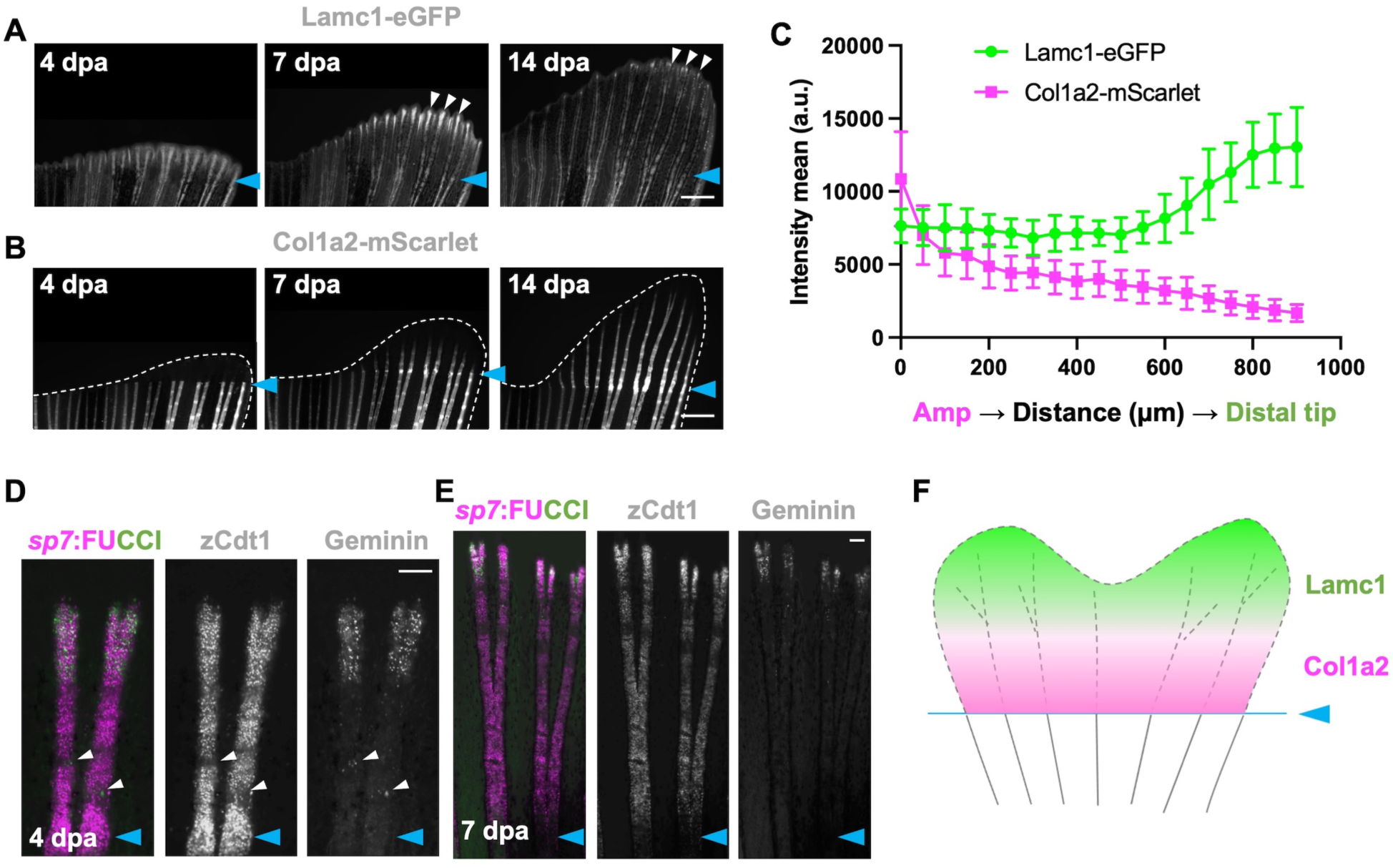
Spatiotemporal deposition of ECM components during fin regeneration indicates a Col1a2-enriched foundation. (A) Regenerating *lamc1^eGFP^* 4, 7, and 14 dpa tail fins. White arrows indicate tip of regenerating tissue. Cyan arrows label amputation plane. (B) Regenerating *col1a2^mScarlet^* 4, 7, and 14 dpa tail fins. Cyan arrows label amputation plane. (C) Quantification results of Lamc1-eGFP and Col1a2-mScarlet fluorescence from amputation plane to distal tips at 7 dpa. Data are shown as mean ± S.D. (D, E) Regenerating *sp7*:FUCCI fin rays at 4 and 7 dpa, indicating cycling (Geminin) and non-cycling (zCdt1) osteoblasts. Arrowheads indicate signals at joints. Cyan arrows label amputation plane. (F) Cartoon indicating spatiotemporal deposition of ECM components during fin regeneration. Scale bars are 500 µm (A-B), 100 µm (D-E)

We were intrigued by the retention of Col1a2-mScarlet localization at the amputation plane -- the junction between newly regenerating tissue and existing tissue. This localization during fin regeneration suggested a potential functional role during regeneration in securing the base or “foundation” of the regenerate. We assessed osteoblast cycling during regeneration by imaging *sp7*:FUCCI fins, harboring reporters that mark cycling (Geminin+) and resting (Cdt+) phases. While we found a concentration of Geminin+ cells at distal areas of growth and at areas of future joints, we did not see measurably higher activity at the base of the regenerate where Col1a2 accumulation occurred (Fig. 4D-F).

To identify molecular changes at the amputation site, we performed bulk RNA-seq from samples collected from the amputation site of regenerating fins at 4, 7, 14, and 21 dpa compared these with 0 dpa samples (Fig. 5A). We found ECM organization and cartilage development genes are enriched at the amputation site, including *col1a2*, *col11a2* (collagen, type XI, alpha 2), *ccn4b* (cellular communication network factor 4b) and chromosome organization genes like *mcm2* (minichromosome maintenance complex component 2) as well as skeletal development genes such as *krt93* (keratin 93) and *mfap2* (microfibril associated protein 2) (Fig. 5B, C). We also observed various ECM genes and matrix metalloproteinase genes such as *mmp9* and *mmp14b* enriched at amputation site compared to samples of tissue from the same region from uninjured fins. We then assessed molecular differences between 7 dpa tissue at the amputation site and the 1 dpa early regenerate, based on published 1 dpa bulk RNA-seq datasets (Kang et al., 2016; Thompson et al., 2020). DNA replication and cell cycle related genes like MCM complex components such as *mcm2*, *mcm3*, *mcm4*, *mcm5* and *mcm6*, Origin recognition complex subunit such as *orc3*, *orc5*, *orc6* and *ccne2* (cyclin E2), *cdk2* (cyclin dependent kinase 2), *cdc45* (cell division cycle 45) and *pola1* (DNA polymerase alpha 1) were highly enriched both in blastema and amputation site (Fig. 5D, E). Notably, collagen genes such as *col1a2* and *col1a1* and collagen metabolism-related genes such as *mmp2*, *mmp9* and *mmp14*, which break down and restructure ECM, accumulated at the amputation site exclusively and not in 1 dpa regenerates (Fig. 5D, E). Thus, the regenerating fin maintains a region at its base that is transcriptionally distinct from the more experimentally targeted distal regenerating portions. The program of this foundation includes long-term deposition of collagen, an event that likely stabilizes the actively growing and maturing fin rays and provides relevant mechanical properties and structural integrity.

**Figure 5.**
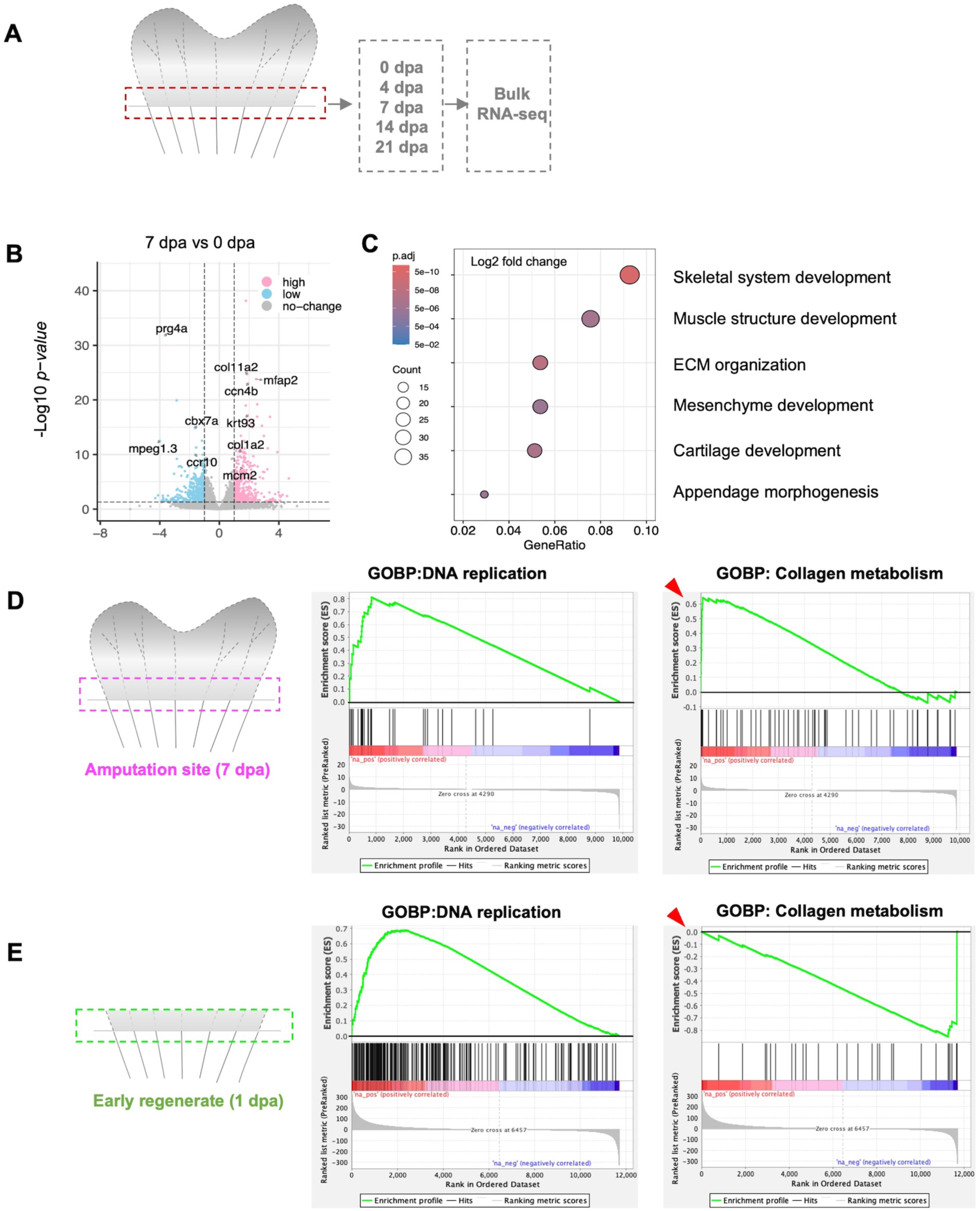
Foundation transcriptome is distinct from uninjured and early regenerating regions. (A) Description of RNA sequencing experiment. Two segments of fin tissue above and below amputation plane were collected from each time point at 0, 4, 7, 14, and 21 dpa. Total RNA was collected for library preparation and sequencing. (B) Volcano plot for gene expression changes, comparing 0 dpa with the 7 dpa amputation site (log2 fold change). (C) GO enrichment terms for gene expression at the 7 dpa amputation site. (D, E) GSEA (Gene Set Enrichment Analysis) analysis of RNA sequencing data for 7 dpa amputation site as compared to the 1 dpa early fin regenerate, comparing gene expression enrichment for DNA replication genes, and collagen-metabolic and catabolic genes. The Y axis is the enrichment score (ES). The black vertical line in the middle panel represents gene hits, indicating where genes from the specific gene set appear within the ranked list. Red color at the bottom of the middle panel indicates high expression in the first condition (Amputation site or 1 dpa), and blue color indicate the high expression in the second condition (similar region from uninjured fins). Red arrowheads highlight differences in y-axis values representing enrichment scores.

### Conclusions

Here we established ECM resources for zebrafish development and tissue regeneration by employing C-terminal in-fusion tagging to visualize endogenous ECM protein dynamics. We detected localization of selected Lamc1, Col1a2 and Tgfbi ECM proteins in various tissues during development and regeneration, and we measured the mobility of ECM proteins in FRAP assays. Finally, the localization of Col1a2 revealed properties of the foundation of regenerating fins with different molecular properties than the distal tip or uninjured regions, highlighting an important, active region of a regenerating appendage that has received little research attention. Future work can expand this toolset to additional ECM proteins, assess regenerative foundations in other models for appendage regeneration, and establish mechanistically how collagens and other proteins contribute to the stability at the base of the regenerating appendage.

## MATERIALS AND METHODS

### Zebrafish

Zebrafish of the Ekkwill strain background were maintained at 28.5 °C with a 14:10 hour light:dark cycle and were used according to protocols approved by Institutional Animal Care and Use Committee (IACUC) guidelines at Duke University or University of Wisconsin-Madison. Adult fish between 6 and 12 months old were used for injury and regeneration experiments. Larvae from 2 to 6 dpf were used for imaging. For larval FRAP experiments, *lamb1a* mutants were maintained at 28°C until 5 dpf, then shifted to a restrictive temperature of 33°C for 24 hours for further analysis. Strains generated in this study are: *lamc1^eGFP^* (allele *pd410*), *col1a2^mScarlet^* (allele *pd411*), *tgbfi^mScarlet^* (allele *pd412*). Other strains used in this study were: *TgBAC(tcf21:DsRed2)^pd37^* (Kikuchi et al., 2011). *sp7:FUCCI* (*Tg(osx:mCherry-zCdt1)^pd270^; Tg(osx:Venus-hGeminin)^pd271^)* (Cox et al., 2018), and *lamb1a^pd110^*(Chen et al., 2015). Larvae were anesthetized with 0.4 mg/ml Tricaine (MS-222; Sigma, A5040) during handling. Adult fish were anesthetized in 0.075% 2-phenoxyenthonal (77699, Millipore Sigma) for surgery. Ventricular resection injuries were performed as previously described (Poss et al., 2002). Zebrafish spinal cord injuries were performed as previously described (Mokalled et al., 2016).

### Generation of C-terminal PCR donors

C-terminal knock-in constructs were gifts from Michel Bagnat’s lab at Duke University. In pUC19 vector backbone, this construct contains a multiple cloning site (MCS), a fluorescent protein (eGFP or mScarlet) coding sequence, stop codon, another MCS, and ubiquitin B (ubb) poly(A) sequence. To modify it to an in-fusion backbone, a flexible linker (aa: GGSGGTGGSGGT) was added between the gene of interest and fluorescent protein(Chen et al., 2013). Gene fragments of partial intron sequence and the last exon from the gene of interest were inserted into the modified pUC19 vector. PCR donors for injection were purified using a GeneJET PCR Purification Kit (Thermo Scientific, K0702). The final products were eluted by nuclease free water and stored at -20°C before injection. Primers for DNA donor: lamc1-forward: 5’-ccttccaagcactgcattcaaattt-3’; lamc1-reverse: 5’-tggccgctctagagacggcgtgtta-3’; col1a2-forward: 5’-aatgtaccaaatttggtttaatacc-3’; col1a2-reverse: 5’-tttgaaacagactgggccaatgtcc-3’; tgfbi-forward: 5’-ggggtgaactcaattatgctgagca-3’; tgfbi-reverse: 5’-ctggacacgagacaatgttctgctg-3’.

### Guide RNA (gRNA) design

gRNA target sites were identified by CHOPCHOP (http://chopchop.cbu.uib.no)(Labun et al., 2016; Labun et al., 2019; Montague et al., 2014). crRNAs and tracrRNAs were purchased from IDT (https://www.idtdna.com/) . gRNAs used in this study were: lamc1-gRNA: 5’-TACTCACTATTATATAGTCG<u>GGG</u>-3’; col1a2-gRNA: 5’-TTCCTTTAGCACACTTAGCA<u>TGG</u>-3’; tgfbi-gRNA:5’-GACCAAATATTGCCAATCGA<u>TGG</u>-3’. Two gRNAs were designed for each gene.

gRNAs were diluted with nuclease-free duplex buffer and stored at -80°C. To test the gRNA efficiency, every gRNA was injected with tracrRNA and NLS-Cas9 (IDT, 10007807) into 100 embryos. 16 injected embryos and 16 uninjected sibling embryos were chosen to prepare genomic DNA, and PCR products from injected and sibling embryos were amplified and annealed. T7EI (T7 Endonuclease I, NEB, M0302) digestion was used to confirm the efficiency of gRNA cutting. Those gRNAs with higher cutting efficiency were used for knock-in establishment.

### Fluorescence Recovery after Photobleaching (FRAP) Analysis

Zebrafish larvae were anesthetized by 0.01% Tricaine and embedded in 1% low-melt agarose on a standard agarose plate. After low-melt agarose solidified, egg water with 0.01% Tricaine was added to dish. FRAP analysis of Lamc1-eGFP, Col1a2-mScarlet and Tgfbi-mScarlet were performed using a Zeiss 880 confocal microscope, with using a 488 nm or 568nm laser and a 20x water immersion lens. Higher laser power was used for photobleaching with 300 repeated cycles. For each experiment, imaging before bleaching was acquired. Images were collected every 10 second for myosepta, and every 30 minutes for fin fold.

### Immunohistochemistry

Zebrafish heart, spinal cord, fin or kidney were fixed in 4% paraformaldehyde (PFA) overnight at 4°C. Heart, spinal cord or kidney were suspended in 30% sucrose and frozen, and 10 μm (heart) or 16 μm cryosections (fin, spinal cord and kidney) were used for immunostaining. Fins were embedded into 1% low melting agar before sectioning to retain tissue structure. Tissue slides were treated by cold methanol for 10 minutes before immunostaining. Slides were dried in room temperature on 37°C for 10 minutes and circled with a hydrophobic pen. Slides were rinsed 2 times in PBST (phosphate buffered saline with 0.1% Tween-20) and blocked with NCS-PBT (10% NCS, heat inactivated newborn calf serum, 90% PBST, 1% DMSO and 2% horse serum) for 1 hour. Primary antibody was applied overnight at 4°C. Slides were washed 4 times for 5 minutes in PBST,then secondary antibody was incubated at 4°C overnight. Slides were rinsed 4 times for 5 minutes in PBST, then treated with Vectashield mounting medium (H-1900, Vector) and DAPI (1:1000, D3571, Thermo fisher). Primary antibodies used in this study were: anti-GFP (1:1000, SKU: GFP-1020, Antibodies inc), anti-RFP(1:500, 5f8, Proteintech), anti-Fibronectin (1:300, F3648, Millipore Sigma), anti-GFAP(Glial Fibrillary Acidic Protein) (1:500, G3893, Millipore Sigma). Secondary antibodies Alexa Fluor 488, Alexa Fluor 567, and Alexa Fluor 633 goat anti-rabbit/mouse/rat/chicken (1:250-1:500, Invitrogen) used in this study.

### RNA sequencing

For caudal fin amputations, fish were anesthetized in 0.075% 2-phenoxyethanol in fish water and fins were amputated with razor blades. 2-3 segment tissue at amputation plane in adult fish were collected after fin amputation at 4, 7, 14, and 21 dpa. Total RNA from 10 pooled fins from three biological replicates were extracted using TRIzol reagent (T9424, Sigma Aldrich) following manufacturer’s instructions. Total RNA was treated with DNase I to clean genomic DNA and purified with Direct-zol RNA MiniPrep kit (R2052, Zymo Research), reverse transcription and preamplification PCR, for cDNA preparation. cDNA purification was performed by adding 0.6x magnetic SPRI beads and washed with freshly prepared 80% ethanol. Then checked cDNA preamplification library content by Qubit. Then Tn5 tagmentation were performed then DNA were purified by DNA Clean & Concentrator-5 kit (Zymo Research, 11-303). Eluted tagmented libraries were loaded into 2% agarose gel and gel size between 300-800 bp were collected, then purified gel by QIAquick Gel extraction kit (Qiagen, 28704). For the final library clean-up, all the tagmented cDNA was pooled together in one sample tube. Libraries were submitted to BGI (Beijing Genomics Institute) and sequenced. RNA-seq reads were trimmed by Trim Galore (v 0.6.7, with cutadapt v 3.5) and counted by salmon (v 1.4.0) by supplying the UCSC danRer11 annotation(Patro et al., 2017). Bioconductor package DESeq2 (v 1.34.0) was used for differential expressions (DE) analysis(Love et al., 2014).

### Datasets

The accession number for the new sequencing data reported in this paper is GEO (Gene Expression Omnibus) GSE291478. We also analyzed the datasets GSE76564 (Kang et al., 2016), GSE146960 (Thompson et al., 2020), GSE75894 (Kang et al., 2016), and GSE193503 (Cigliola et al., 2023).

## Acknowledgments

We thank the Duke Zebrafish Core and Jim Burris and Mackenzie Klemek at Morgridge Institute for Research for zebrafish care; C.-H. Chen and J. Cao for helpful comments on the manuscript. The authors acknowledge funding from NIH (R35 HL150713 and R01 HD105033 to K.D.P) and from Pew Charitable Trusts to D.R.S. and K.D.P.

## SUPPLEMENTARY FIGURES

**Figure S1.**
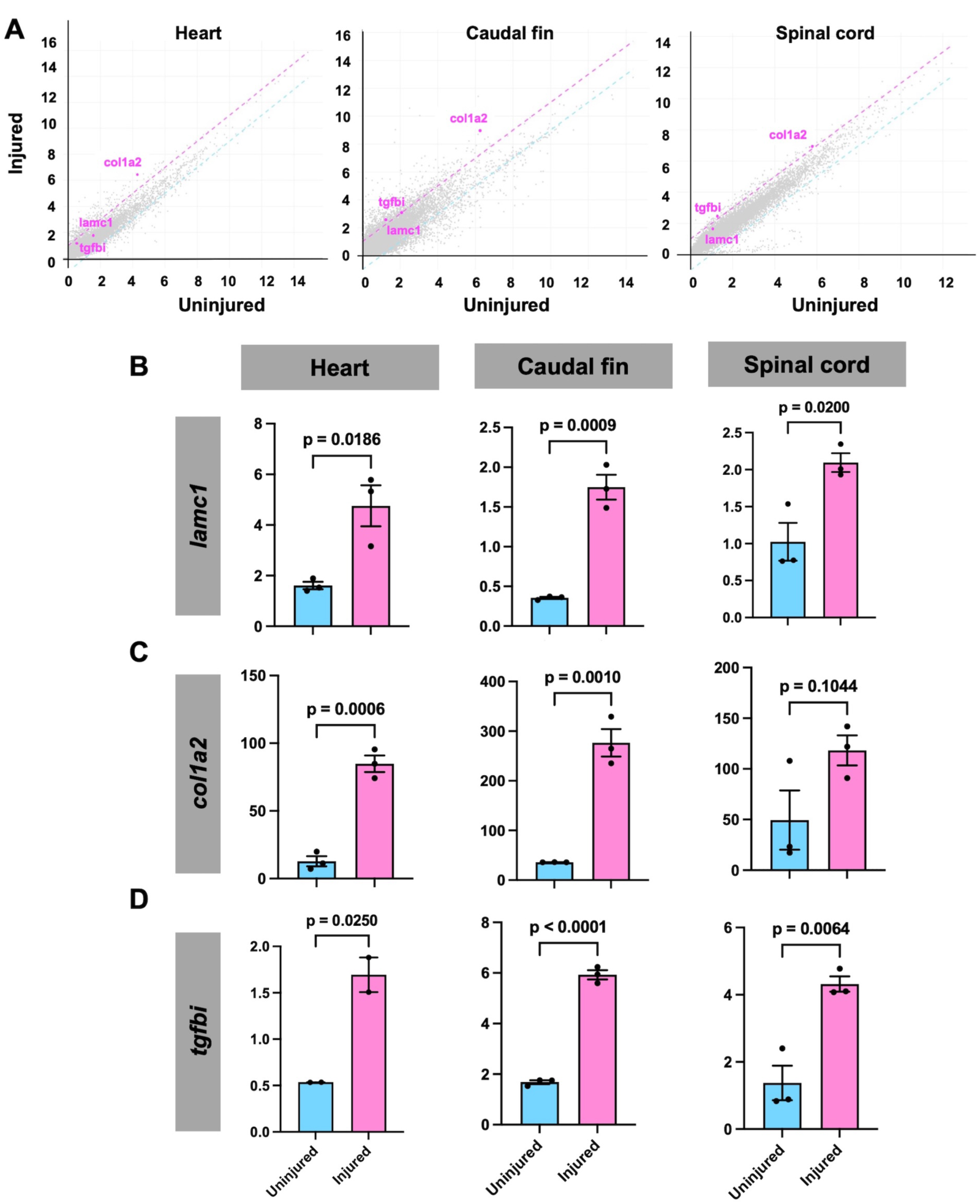
Expression levels of *lamc1*, *col1a2* and *tgfbi* during tissue regeneration. (A) XY scatter plots of differentially regulated genes during heart, fin, and spinal cord regeneration. Magenta lines indicate thresholds for significantly increased RNA levels, and cyan lines indicate thresholds for significantly decreased RNA levels. Cutoff fold changes: log2>1 for increasing and log2<-1 for decreasing. (B-D) RPKM (Reads Per Kilobase per Million mapped reads) of mRNA levels of *lamc1*, *col1a2* and *tgfbi* in uninjured and regenerating caudal fins (4 dpa), cardiac ventricles (7 days post ablation), and spinal cords (7 days post transection). Student’s *t*-test.

**Figure S2.**
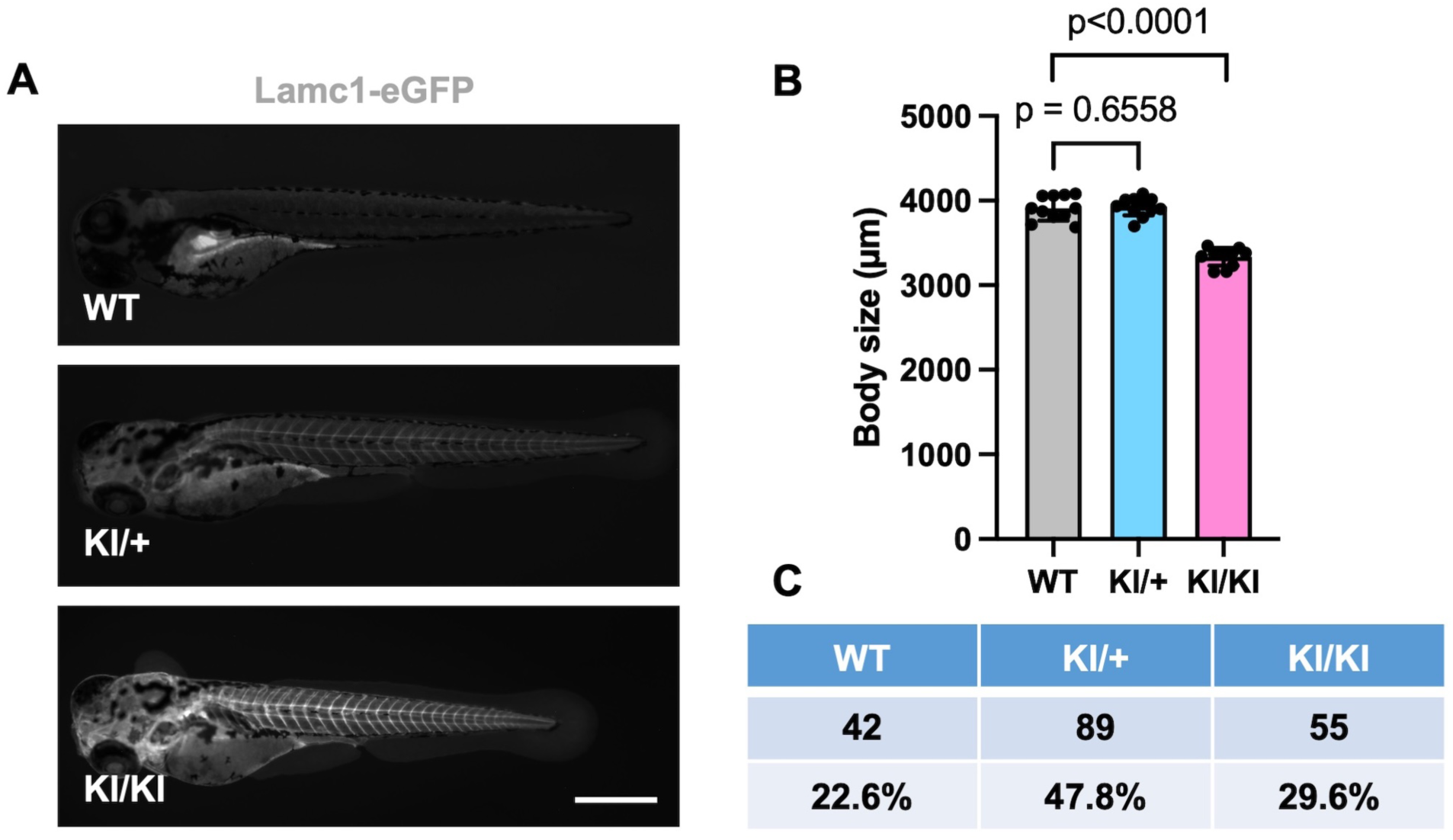
Images and survival of homozygous *lamc1^eGFP^* larvae. (A) Representative pictures for WT (wild-types), *lamc1^eGFP^*heterozygotes and *lamc1^eGFP^* homozygous larvae at 4 dpf. (B) Body lengths of wild-types, *lamc1^eGFP^* heterozygotes, and *lamc1^eGFP^* homozygous larvae at 4 dpf. n = 10 from each group. Student’s *t*-test. (C) Ratios of wild-types (EK), *lamc1^eGFP^* heterozygotes, and *lamc1^eGFP^* homozygotes at 4 dpf. Scale bar is 500 µm in (A).

**Figure S3.**
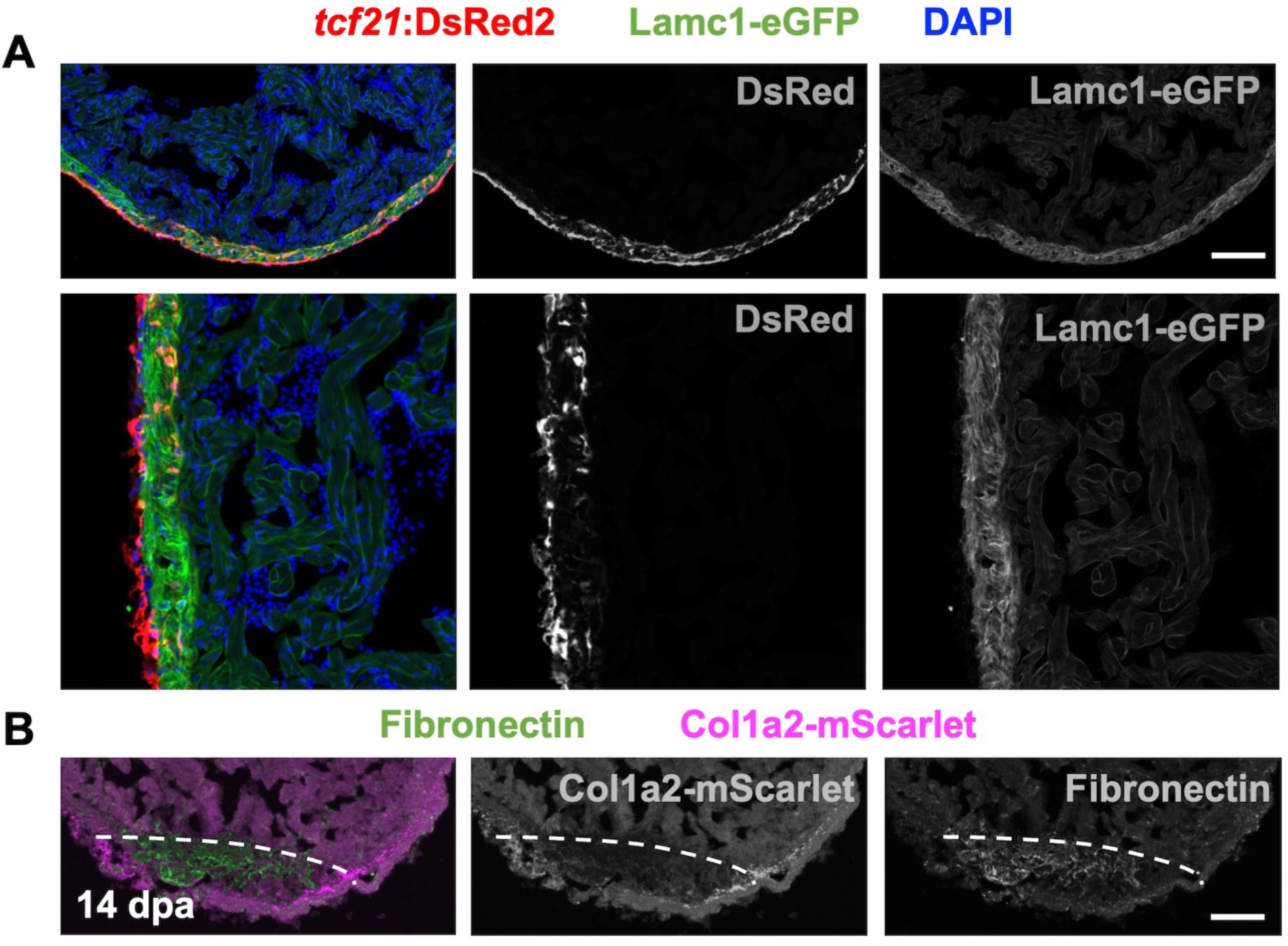
Cardiac expression of ECM components. (A) Confocal image of Lamc1-eGFP in section from uninjured adult ventricle, co-stained for epicardial and epicardial-derived cells (*tcf21*:DsRed2) and nuclei DAPI (blue). (B) Confocal image of section of 14 dpa ventricle, co-stained for Fibronectin and Col1a2-mScarlet. Scale bar is 100 µm (A, B).

**Figure S4.**
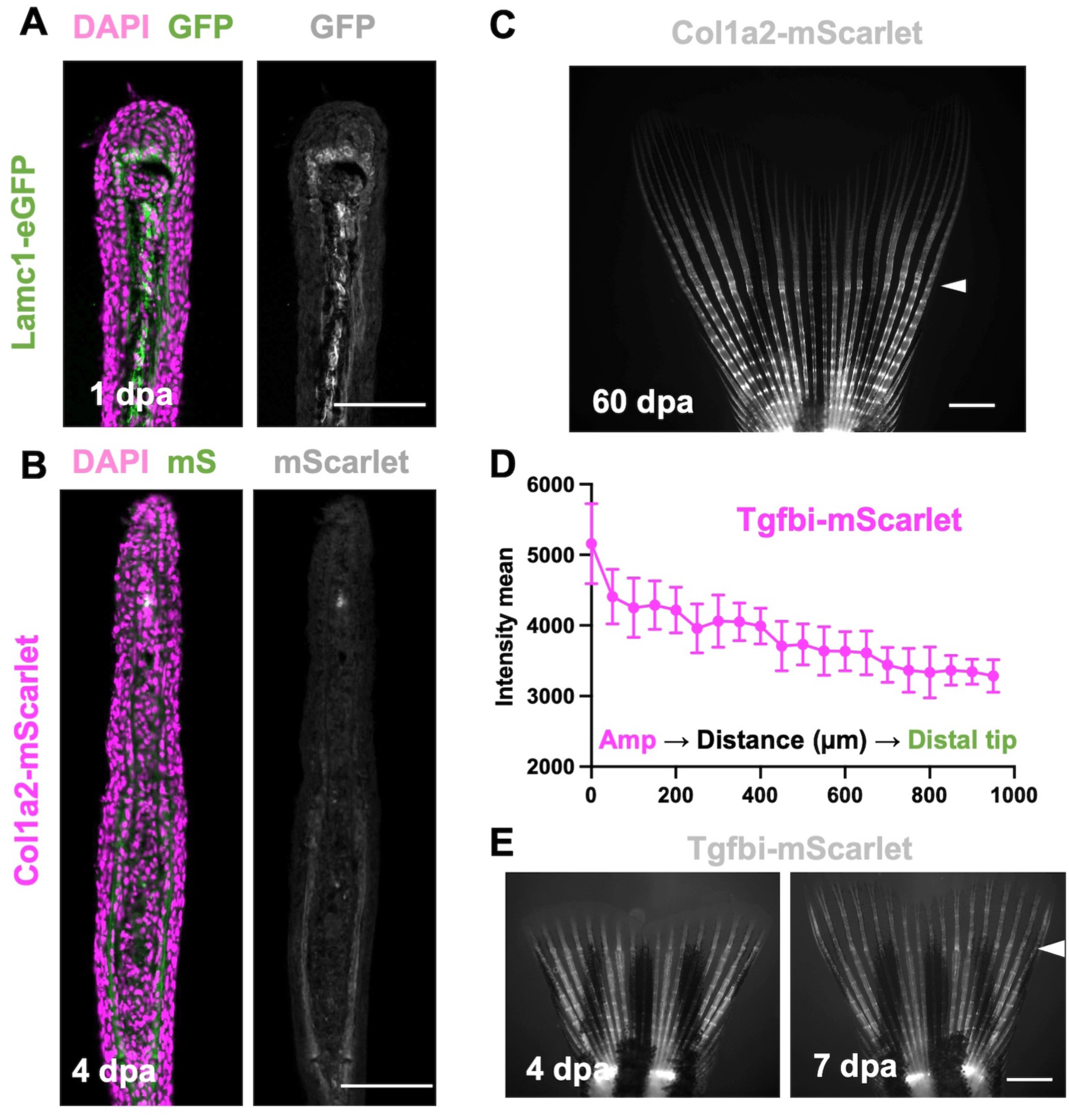
ECM components during fin regeneration. (A, B) Section of fin regenerates indicating Lamc1-eGFP and Col1a2-mScarlet at 4 dpa. (C) Image of regenerating fin at 60 dpa, indicating persistent Col1a2-mScarlet localization at the amputation site. (D) Quantification of Tgfbi-mScarlet fluorescence in regions of 7 dpa regenerating fins from the amputation plane (Amp) to the distal tip. Data are mean ± S.D. n = 5, 4 fin rays were quantified from each fish. (F) Representative images of regenerating *tgfbi^mScarlet^* fins at 4 and 7 dpa. Scale bars are 100 µm in (A, B), 1 mm in (C, E).

